# Mechanical instabilities of aorta drive blood stem cell production: a live study

**DOI:** 10.1101/509190

**Authors:** Nausicaa Poullet, Ivan Golushko, Vladimir Lorman, Jana Travnickova, Dmitryi Chalin, Sergei Rochal, Andrea Parmeggiani, Karima Kissa

## Abstract

During embryogenesis of all vertebrates, haematopoietic stem/progenitor cells (HSPCs) extrude from the aorta by a complex process named Endothelial-to-Haematopoietic Transition (EHT). HSPCs will then colonize haematopoietic organs allowing haematopoiesis throughout adult life. The mechanism underlying EHT including the role of each aortic endothelial cell within the global aorta dynamics remains unknown. In the present study, we show for the first time that EHT involves the remodelling of individual cells within a collective migration of endothelial cells which is tightly orchestrated, resulting in HSPCs extrusion in the sub-aortic space without compromising aorta integrity. By performing a cross-disciplinary study which combines high resolution 4D imaging and theoretical analysis based on the concepts of classical mechanics, we propose that this complex developmental process is dependent on mechanical instabilities of the aorta preparing and facilitating the extrusion of HSPCs.

## Introduction

Transplantation of human blood cells is essential to regularly save lives on a large-scale. However, this method requires an exogeneous allogenic source to avoid histoincompatibility as well as graft versus host disease associated problems. Currently, haematopoietic stem/progenitor cells (HSPCs) can only be produced *in vitro* by methods involving genetic cellular reprogramming. Different groups succeeded in it, however these approaches remain scientifically and technically challenging ^1–4^. Moreover, the presence of transgenes in the genome of reprogrammed human HSPCs represents an important clinical risk ^5^.

In order to develop new methods to generate HSPCs *in vitro* and control their fate after transplantation, we need to further deepen our knowledge on blood cells ontogenesis at tissue and organism levels, considering also novel features like biomechanical forces experienced by HSPCs in physiological conditions. Indeed, the 3D *in vivo* structure of the tissue from which HSPCs are generated is subjected to mechanical forces that influence cellular properties and processes like cell migration, adhesion and polarity ^6,7^ Moreover, mechanical stress is known to have a major impact on gene expression modulation and consequently on developmental processes ^8^, inflammation and cancer ^9^.

In this article, we address for the first time the question of HSPCs production from the aorta in relationship with the growth of the zebrafish embryo, the most widely used animal model to study developmental processes in real time. We discuss and underline that, together with haemodynamic forces, the growth of the whole embryo generate mechanical stresses on the aorta and play a crucial role in blood production.

Previously, we and colleagues have demonstrated that HSPCs emerge in the Aorta Gonad Mesonophros (AGM) region ^10–13^, from the ventral wall of the dorsal aorta (DA) ^14–16^. We named this process Endothelial-to-Haematopoietic Transition or EHT ^14^ In zebrafish, EHT takes place during a specific time window between 30 and 65 hours post fertilization (h.p.f.)^12,14,17^ Systematic tracking of aortic endothelial cells (EC) in live embryos showed that HSPCs emerge from the aortic ventral floor through a process that involves a strong shape change followed by the egress of single cells from the aortic ventral wall into the sub-aortic space ^14^ Moreover, we and colleagues observed that the extrusion of HSPCs was aborted when the *runx1* transcription factor essential for HPSCs emergence was inhibited^14,18^. Surprisingly, this inhibition affected neither aorta formation nor any events preceding HSPCs extrusion, such as aorta radius dilation and contraction. A more recent study has then put in evidence with wealth of details the cytoskeletal processes occurring in the single cell dynamics during the HSPC egress from the aorta^19^.

In the present study, with the help of 4D confocal microscopy, we follow and quantify at tissue level, the whole aorta behaviour as well as the one of its cells throughout EHT between 24 and 72 h.p.f. We put in evidence important structural changes of the aorta and the collective migration of its lateral cells down to the aorta floor prior to HSPC egress. These phenomena result in a global aorta remodelling in terms of number and localisation of endothelial cells, thus showing an ongoing global cellular reorganisation of the tissue that assures aorta integrity during EHT. We then analyse the role of the actin cytoskeleton both in emerging and neighbouring cells during the HSPCs extrusion process.

Based on these observations and applying general principles of mechanics to the novel context of EHT, we relate the overall aorta remodelling and EHT egress to mechanical instabilities. This cross-disciplinary analysis indeed reveals that mechanical instabilities, induced by different stresses arising from the inhomogeneous growth of the aorta and its interaction with surrounding growing tissues, play a key non-specific role in HSPCs extrusion. Thus, not only haemodynamic forces, but also stresses induced by the global zebrafish embryonic growth ^17^ are essential for haematopoiesis.

## Results

### EHT is associated with important aorta remodelling

To investigate the mechanisms underlying EHT, we first carried out a 4D confocal microscopy imaging of this process in a physiological context. For that, we image the trunk of zebrafish embryos (Fig. 1a) including diverse structures such as the DA, the cardinal vein (CV), the notochord and the muscles surrounding the DA (Fig. 1a-c) between 30 and 65 h.p.f.

**Figure 1.**
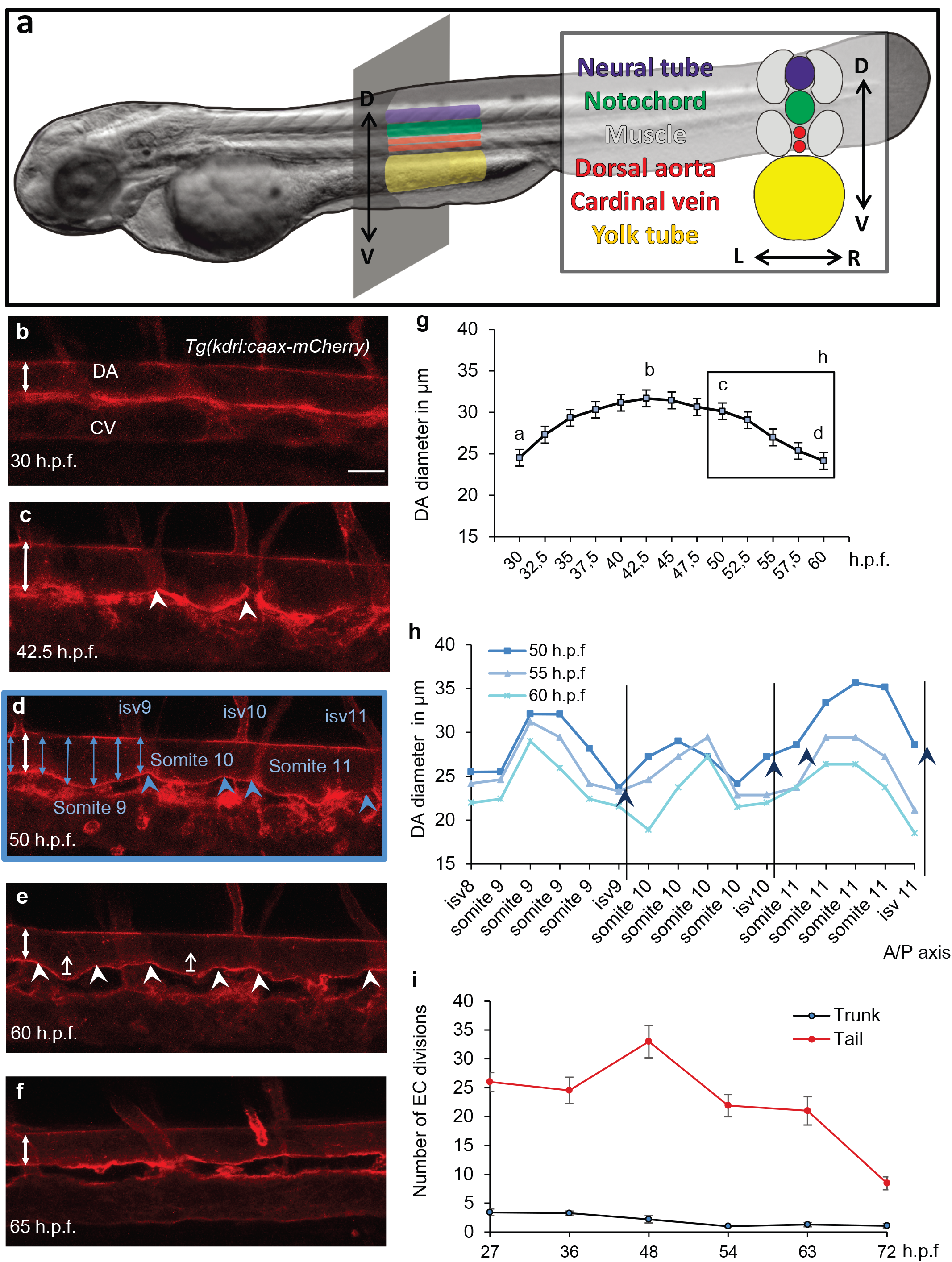
Aorta remodelling during EHT: embryo anatomy and spatiotemporal organisation of aorta diameter. **a**. Drawing showing the AGM localization in the trunk of a 48 h.p.f. zebrafish. Scheme of the AGM in longitudinal or transverse views (grey boxes) consisting of the neural tube, the notochord, the dorsal aorta, the cardinal vein, and the yolk tube. **b-f**. Still frames of time-lapse imaging of *Tg*(*kdrl:caax-mCherry*) embryo from 30-65 h.p.f. Maximum projection from 40 z-stack spaced by 1μm. Double-headed arrows indicate difference of aorta diameter through time. **b-c**. Between 30 and 42.5 h.p.f., DA diameter expands. Arrowheads indicate localization of cells leaving the aorta and forming a local reduction of the diameter. **d**. Image of *Tg*(*kdrl:caax-mCherry*) embryo at 50 h.p.f. to illustrate DA diameter measurements throughout the length of 3 somites as represented in **h. e**. From 45 to 65 h.p.f., DA diameter decreases, corresponding to the peak and then decline of EHT events. Arrows indicate the contraction of the aorta occurring from the floor. **f**. After EHT, at 65 h.p.f, aorta floor becomes flat. **g**. Graph showing aorta diameter variation in time measured every 2.5h in 5 embryos. Letters correspond to the according panels. Error bars represent standard error of the mean. **h**. Graph showing variation of diameter along the antero/posterior axis at 3 time points: 50, 55 and 60 h.p.f. (Black box from g) in the embryo imaged in b-f. Dotted line indicates intersomitic vessels. Blue arrowheads indicate localization of EHT cells leaving the aorta and forming a local reduction of the diameter. **i**. Graph showing the number of EC divisions *(kdrl:mV-zGmn*^+^ cells) occurring between 27 and 72 h.p.f. in the trunk and in the tail region, n=10 cells. Note that a peak of division is observed in the tail region between 36 and 54 h.p.f. corresponding to the formation of the CHT and its colonisation by HSPC. DA: dorsal aorta; isv: intersomitic vessel; CV: cardinal vein. Scale bar: 25 μm. See also **Supplementary Movie S1**.

The imaging revealed strong changes of the morphology of the whole DA with time (Fig. 1b-f, **Supplementary Movie S1**). From 24 to 42.5 h.p.f., we observed a drastic increase of the aorta diameter from 24±0.8 μm to 32±0.9 μm (Fig. 1c, 1g) followed by the emergence of a pattern with alternating thinner and thicker diameter regions (Fig. 1c-d, 1g-h) with a relative amplitude variation ranging from 17% to 33% the average aorta diameter.

At 42.5 h.p.f., the average diameter of the DA starts to decrease (Fig. 1f, 1g) and at 65 h.p.f., the aorta original cylindrical shape and diameter are restored which corresponds to the end of EHT (Fig. 1d-f, 1g). To determine if there is a causal link between the change in aorta diameter and cell extrusion, we followed the behaviour of individual cells leaving the DA (Fig. 1e-f, arrowheads correspond to regions where HSPCs emerged). Our observations showed that interestingly, HSPC extrusion rate peaks between 42.5 and 52 h.p.f. precisely when the aorta diameter starts to decrease.

### Aorta dilation is not associated with cell mitosis

To further assess EC behaviour during aorta dilation and HSPC extrusion, we looked at EC division and whether an increase of cell number can explain vessel expansion or compensate for HSPC emerging from the aorta floor. We used a zebrafish transgenic line that expresses a marker of the S/G2/M phase of the cell cycle, mVenus-zGeminin ^20^. This marker was specifically expressed under the *kdrl* promoter, allowing visualization of ECs during mitosis ^20^. We quantified the number of EC divisions in the trunk and in the tail between 27 and 72 h.p.f. We found that many division events occur in the tail region (33.0±2.8 EC divisions at 48 h.p.f), allowing development of the Caudal Haematopoietic Tissue (CHT) (Fig. 1i). In contrast, at the peak of EHT (at around 48 h.p.f.), we quantified a mean of 2.2±0.6 ECs dividing in the *whole* AGM area (Fig. 1i), far under what would have been needed to compensate for the estimated 25 HSPCs emerging from the aorta during the whole AGM ^14^ Overall, between 27 and 72 h.p.f., we observed only a few EC divisions in the trunk region: the total number of cells in this area essentially decreases throughout the EHT process. Thus, additional surface area, for the aorta dilation and later for integrity maintenance during HSPC extrusion, should be provided exclusively by cell deformation.

### HSPCs migrate collectively from the sides towards the ventral part of the aorta

To quantify the precise contribution of each EC to the EHT process, we followed the fate and behaviour of individual cells between 30 and 60 h.p.f. in the whole trunk region of the DA. To do so, we used the double transgenic line *kdrl:nls-GFP/kdrl:caax-mCherry*, allowing respective visualization of nucleus and membrane of ECs through time. We discovered that ECs undergo substantial rearrangements in the trunk region between 30 and 45 h.p.f. (Fig. 2a-b). Interestingly, during this time window ECs migrate massively from the lateral side of the DA to the floor (Fig. 2a-h with arrows). The number of cells in the DA floor peaks at about 40 h.p.f., when cell migration from the sides fully compensate for cell egress, and then steadily declines afterwards (Fig. 2i) as lateral cells stop migrating towards the DA floor. The decline is due to the extrusion of cells from the aorta floor into the sub-aortic space. The number of ECs forming the dorsal aorta in the trunk region therefore strongly decreases from the start to the end of the process (from 12.1±0.9 cells/somite at 30 h.p.f. to 7.9±0.3 cells/somite at 60 h.p.f., p-value=0.009) (Fig. 2a-c, 2i-j). Remarkably, the number of ECs localized in the DA roof (facing the notochord) is essentially constant during the whole process (Fig. 2i).

**Figure 2.**
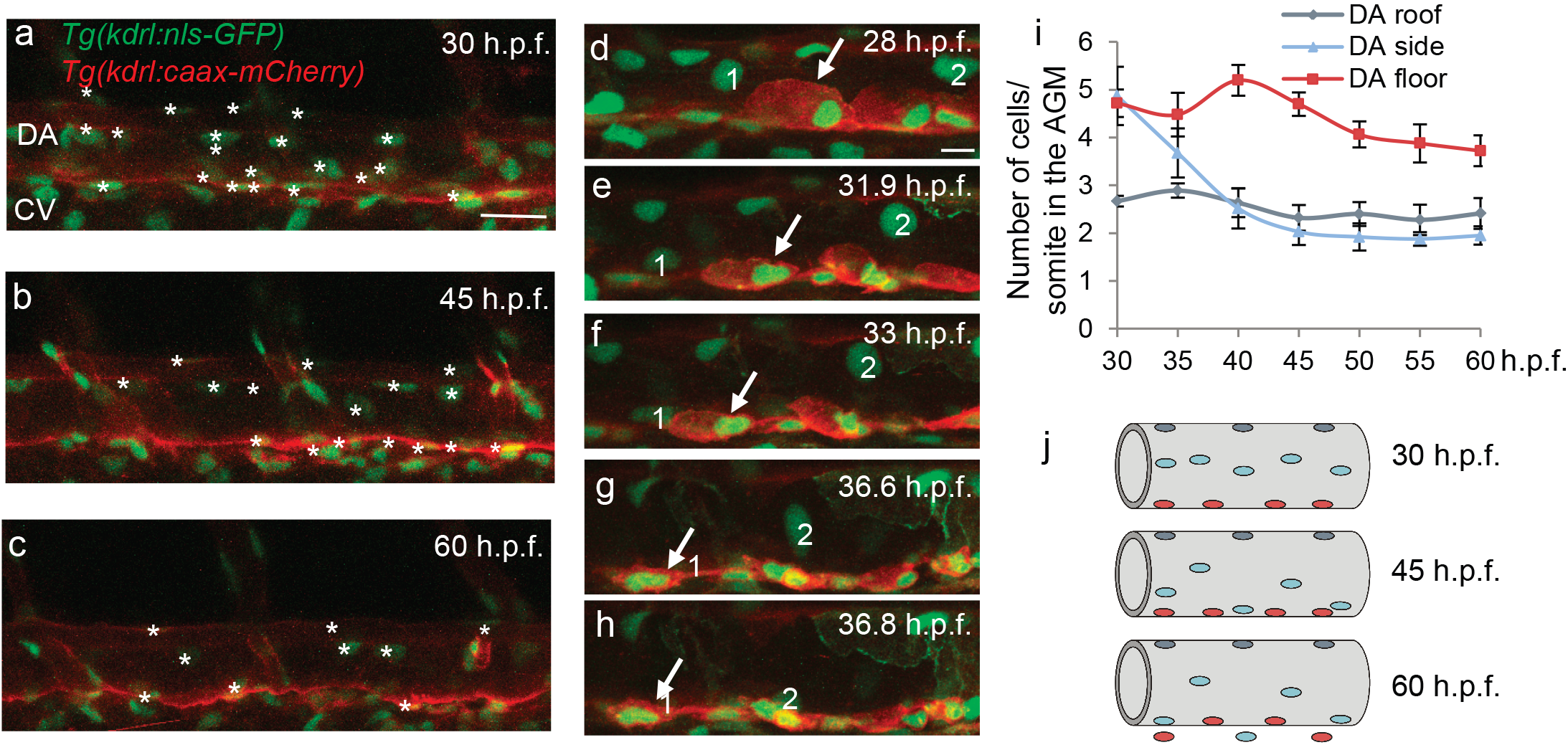
Cells undergoing EHT are recruited from the side of the aorta and migrate to its floor prior to extrusion. **a-c**. From 30 to 60 h.p.f., still frames of time-lapse imaging of *Tg*(*kdrl:ns-GFP)/Tg(kdrl:caax-mCherry*) embryo. Maximum projection from 40 z-stack spaced by 1μm. Stars indicate the nucleus of the ECs in the dorsal aorta in the 2 central somites of the image. **d-h**. Still frames of time-lapse imaging of *Tg*(*kdrl:nls-GFP)/Tg(kdrl:caax-mCherry*) embryo. Maximum projection from 40 z-stack spaced by 0.6μm. Cells migrating from the side of the aorta toward the floor are numbered 1 and 2 in white. **i**. Graph showing the number of nuclei counted in 3 different zones of the dorsal aorta: roof, side and floor, n=5 cells. **j**. Schematic representation of cell rearrangement occurring from 30 to 60 h.p.f. Colour coding of the cells corresponds to the graph in **i**. DA: dorsal aorta; CV: cardinal vein. Scale bar: 25 μm (**a-c**), 10μm (**d-h**).

### HSPCs extrusion requires collective EC morphology changes

In order to determine the contribution of neighbouring ECs to the extrusion of HSPCs, we compared the morphology of the emerging cells versus the morphology of their neighbours during EHT. Imaging of the double transgenic line *kdrl:utrophin-CH-GFP/kdr:nls-GFP* allowed to visualize the boundaries of each individual EC together with its nucleus. Quantification of the cell area between 40 and 60 h.p.f. showed that both emerging cells and their neighbours undergo drastic and rapid morphology changes (Fig. 3). The area of cells prior to their extrusion significantly reduces and shape changes from flat endothelial to a round plate-like cell (−69%±5% decrease in cell area) (Fig. 3a-b, 3e-g). In contrast, neighbouring lateral cells increase their area comparably (+60%±9% increase in cell area) (Fig. 3c-d, 3e-f and 3h). These data confirm that in the absence of divisions in the trunk region, the loss of cells through EHT is mainly compensated by the deformation of the lateral ECs thus assuring global aorta integrity.

**Figure 3.**
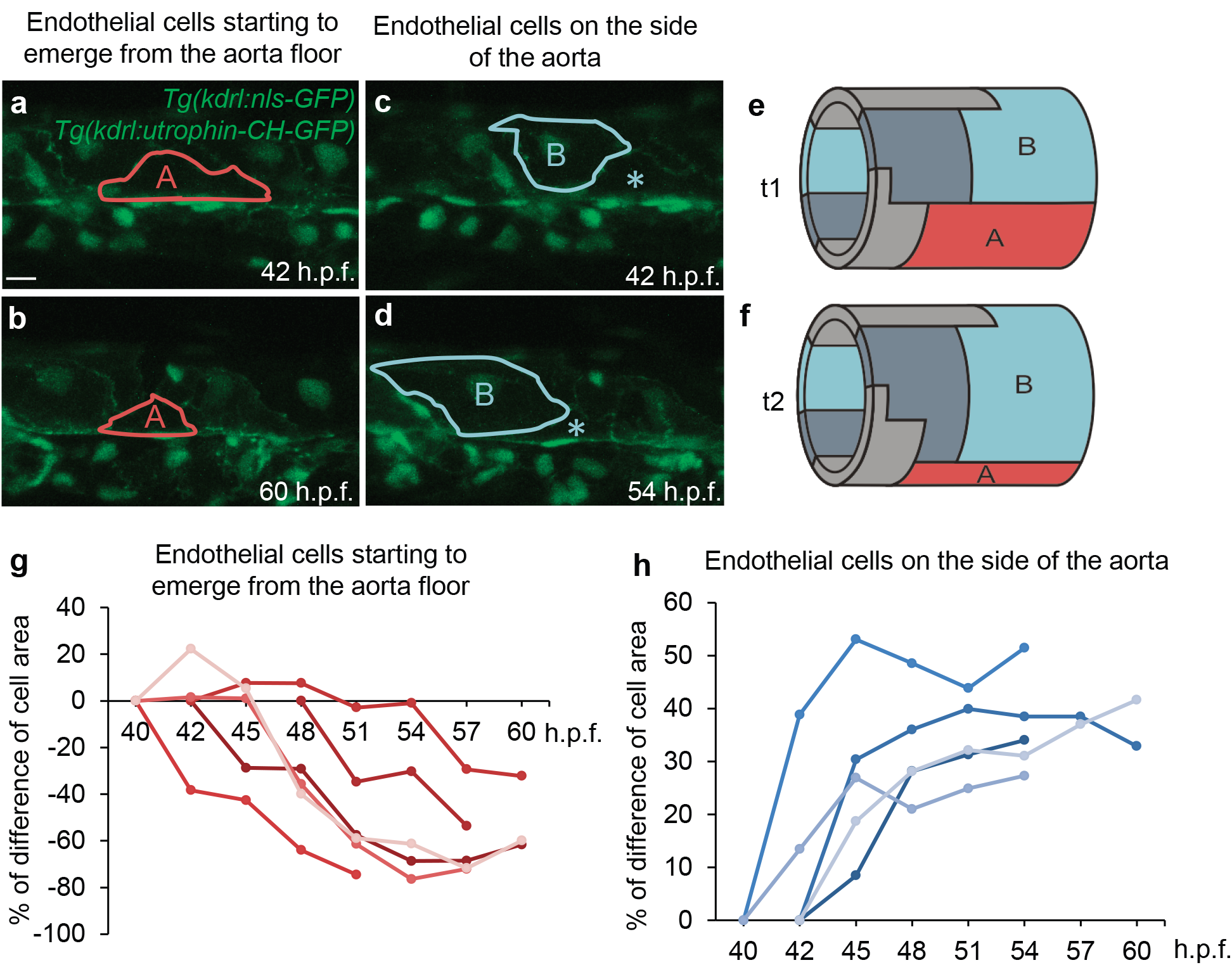
Endothelial cells adopt collaborative behaviour during EHT: cellular contractions and extensions to maintain aorta integrity. **a-d**. Still frames of time-lapse imaging of *Tg*(*kdrl:nls-GFP)/Tg(kdrl:utrophin-CH-GFP*) embryo between 40 and 60 h.p.f. Maximum projection from 40 z-stack spaced by 0.6μm. Cell A (highlighted in orange) is an endothelial cell located in the floor of the aorta and starting to undergo EHT. Cell A surface reduces importantly between 42 h.p.f. (a) and 60 h.p.f. (b). Cell B (highlighted in blue) is located in the side of the aorta neighbouring a cell undergoing EHT (star). Cell B surface increases between 42 h.p.f. (c) and 54 h.p.f. (d). e-f Schematic representation of EC surfaces and positions at t1 (e) and t2 > t1 (f). Colour-code and letters correspond to the panel a-d. **g**. Graph showing the temporal evolution of EC starting to emerge from the aorta floor in percentage of difference compared to area at 40 or 42 h.p.f. Each line represents the measurements for one cell. 6 ECs starting to emerge from the aorta floor were analysed in 5 different embryos. For details on cell area (μm^2^) calculation, see Material and Methods section. **h**. Graph showing temporal evolution of cell area of lateral EC in percentage of difference compared to area at 40 or 42 h.p.f. Each line represents the measurements for one cell. 5 ECs on the side of the aorta were analysed in 5 different embryos. Scale bar: 10μm.

### Extrusion is finalized by actin ring closure around the emerging HSPC

We then focused on the cytoskeletal activity of ECs to understand the mechanism of this dramatic cell-to-tissue reorganization process. First, we used the *kdrl:utrophin-CH-GFP* line to mark stable F-actin in ECs ^21^. 4D confocal imaging during the period of rearrangement (42-60 h.p.f.) showed that EC junctions are highly dynamic and have tight cell-to-cell membrane boundaries (**Supplementary Movie S2**). Because of the significant vascular cell deformation during EHT, we then analysed the interplay between cytoskeleton activity and HSPCs shape during the extrusion process. We imaged a double transgenic line *kdrl:utrophin-CH-GFP/kdrl:caax-mCherry* showing respectively stable actin filaments and EC membranes. Live imaging revealed that the aorta diameter decreases while HSPC extrusion in the sub-aortic space are driven by the formation and closure of an actin ring surrounding the emerging cells (Fig. 4, **Supplementary Movie S1, Supplementary Movie S3**) as also recently observed by Lancino et al ^19^. Transverse view of the aorta shows that the emerging cell membrane and the actin cytoskeleton co-localize to form a perfect circle that closes as the cell exits the DA floor (Fig. 4a-c **Supplementary Movie S3**). As the *kdrl:utrophin-CH-GFP* line displays a mosaic expression, not all ECs express Utrophin-CH-GFP and we were therefore able to follow EHT in cases where only the neighbouring cells expressed the GFP probe. In this scenario, we still clearly observed the formation (Fig. 4a, 4d) and closure of the actin ring (Fig. 4b-c, e-f), suggesting that the neighbouring cells also actively participate in the ring formation and actin contractile dynamics.

**Figure 4.**
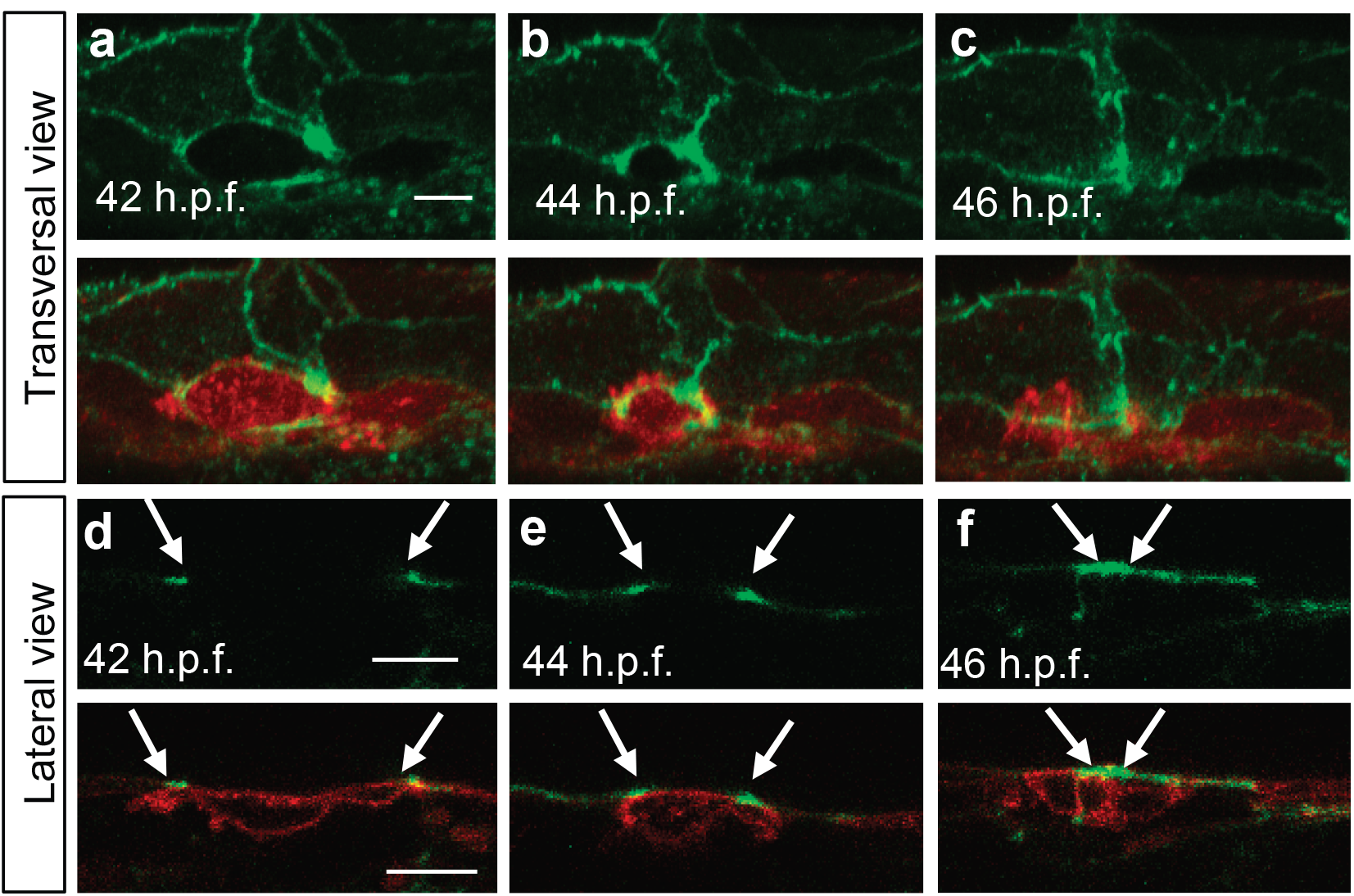
Final contraction in EHT is actin-dependent and coordinated with surrounding cells. **a-c**. Horizontal reconstructed view from a maximum projection of a series of z-stack of a *Tg*(*kdrl:utr-CH-GFP)/Tg(kdrl:caax-mCherry*) embryo during EHT at 42, 44 and 46 h.p.f. Upper panels are *kdrl:utrophin-CH-GFP* alone and lower panels are merged images with *kdrl:caax-mCherry*. Closure of the actin ring by the neighbouring cells is clearly observable in the upper panels. See also **Supplementary Movie S3. d-f**. Single z-stacks of time-lapse imaging of the same embryo as in a-c. Upper panels are *kdrl:utrophin-CH-GFP* alone and middle panels are merged images with *kdrl:caax-mCherry*. Actin ring (arrows) closes around emerging cell (arrowhead). Note that in this case the emerging cell does not express Utrophin-CH-GFP due to mosaic labelling, allowing us to confirm the role of the neighbouring cells in actin ring closure.

In order to visualize all F-actin in the system, we also used a transgenic line expressing Lifeact (a 17-amino-acid peptide that binds to filamentous actin) fused to green fluorescent protein (GFP) ^22^. Vascular expression of Lifeact-GFP was driven by the VE-cadherin promoter and this transgenic line was used together with *kdrl:caax-mCherry* to visualize EC membrane as well. Imaging of *VE-cad: Lifeact-GFP* was consistent with the results obtained with *kdrl:utrophin-CH-GFP* and we could follow the formation and closure of the actin ring during EHT (data not shown).

To confirm the role of actin polymerization in HSPC extrusion, we treated embryos with drugs blocking actin assembly, Latrunculin B or Rac inhibitor NSC23766. In treated embryos, despite the presence of the aorta diameter modulation, the actin cytoskeleton was highly disrupted and the formation of the actin ring surrounding the emerging cells was not observed (**Supplementary Fig. 2**). Interestingly, the formation of typical plate-like cells in the aorta floor preceding HSPCs extrusion was also not observed (data not shown). We also found that in treated embryos some cells undergoing EHT bent toward the DA lumen: 33%±9% of cells emerged toward DA lumen in Latrunculin B-treated embryos (**Supplementary Fig. 2**). These results show the importance of actin polymerization for the acquisition of cell rigidity and the direction of HSPCs extrusion from the aorta in the sub-aortic space.

We further looked at the role of the acto-myosin contraction in HSPC extrusion by subjecting embryos to drugs blocking myosin contraction, Blebbistatin and ROCK inhibitor Y-27632. The action of these drugs was very similar to that of the actin-blocking ones. The actin ring surrounding emerging cells was not observed and HSPC exit took longer with some cells undergoing fragmentation during the process (**Supplementary Fig. 2**).

To quantify the effect of actin polymerization and contraction blocking on haematopoietic organs colonization such as the CHT by HSPCs, we treated *cd41:GFP* embryos, which express GFP in HSPCs and thrombocytes ^23^, with Latrunculin B, Rho-kinase inhibitor Y-27632, Blebbistatin and Rac inhibitor NSC23766. For all drugs, except with Rho-kinase inhibitor Y-27632, we found that the number of HSPCs colonizing the CHT (cd41+ cells) between 52 and 72 h.p.f. was significantly lower in treated embryos compared to control (Supplementary Table 1). Live imaging of EHT reveals that HSPCs bend without achieving complete extrusion. Part of them bents in the wrong direction, i.e. in the aorta lumen, while others cells burst as previously observed after *runx1* gene inactivation ^14^. Interestingly, none of the drugs blocking actin polymerisation or contraction was found to affect EC rearrangement occurring during the earlier stages of EHT (data not shown).

All together these results show that the acto-myosin cytoskeleton plays an essential role mainly in the final step of EHT, precisely to complete the extrusion of HSPCs from the aorta floor to the sub-aortic space.

### Cellular and tissular levels are dynamically coordinated during EHT

Structuring up our observations, the EHT transition is a collective phenomenon organized on both tissular and cellular spatial scales (Fig. 5).

**Figure 5.**
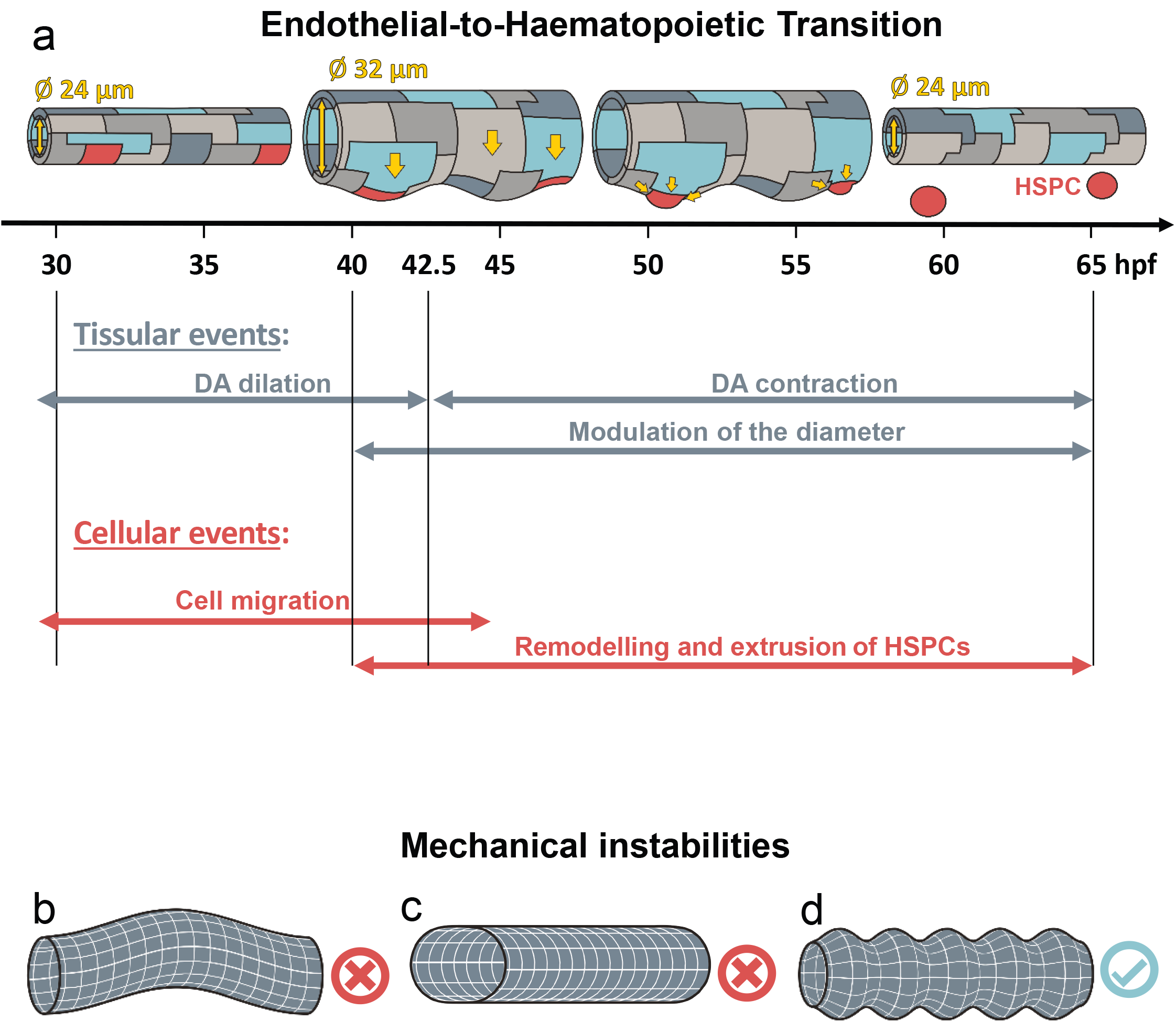
Overall schematic representation of Endothelial-to-Haematopoietic Transition with mechanical instabilities. **a**. The EHT transition starts first by a global aorta dilation (as soon as the heart starts to beat), taking place from 24 to about 42,5 h.p.f., followed by the aorta contraction, occurring from about 42,5 to 65 h.p.f. While the embryo is growing and developing, the first phase of aorta dilation is essentially characterized by cells that start to increase their area, while others, in particular those located on the sides of the aorta, migrate towards the ventral part (aorta floor). During this phase the characteristic modulation of the aorta diameter, the corrugation instability (see main text and below), appears. In the contraction phase, cells localized at the aorta floor undergo a drastic morphological change, shaping from flat ECs to round plate-like cells. As cells round up, they undergo a strong antero-posterior contraction. In parallel, neighbouring ECs compensate for the surface reduction of emerging cells, and eventually for the extrusion of the latter cells from the aorta, by increasing importantly their surface. Finally emerging round cells extrude and individualize from the aorta. This phase coincides with the decrease of the diameter modulation, recovering the initial aorta diameter and its cylindrical geometry. **b-c-d**. The three main mechanical instability modes of a thin cylindrical soft pipe (membrane) under mechanical stresses: **b**. Longitudinal buckling; **c**. Transverse buckling; **d**. Corrugation.

At tissue level, EHT is organized in two phases. In the first phase, from 28 to 42.5 h.p.f. the DA diameter increases (Fig. 1g and 5a). At 40 h.p.f. aorta’s shape of straight cylinder is distorted by an average diameter modulation along the aorta longitudinal direction (Fig. 1h and 5a).

During this first tissular phase, between 30 and 40 h.p.f., and concurrent with the diameter modulation, at cellular level ECs reorganize themselves spatially and cells, localized on the side of the aorta, migrate towards its floor. This cell migration is a quick process, which takes on average 5.9h±0.7h (counted for 18 cells in 15 individuals).

The second tissular phase corresponds to the diameter reduction and the disappearing of the diameter modulation up to the end of EHT (from about 42.5 to 65 h.p.f.). Here, global morphological reorganization of DA is coupled with the dynamics of cells that, once localized at the aorta floor, undergo a drastic morphological change, shaping from a flat EC to a round plate-like cell. As cells round up, they undergo a strong antero-posterior contraction. In parallel, neighbouring ECs increase their area and compensate for the surface reduction of emerging HPSC cells and eventually for their extrusion. The whole cellular dynamics assures aorta integrity while DA diameter decreases. This important morphology and identity tissular and cellular remodelling take on average 9.2h±1.5h (counted for 6 cells in 6 individuals) and occur between 37 and 55 h.p.f. The closure of the actin rings around the emerging cells allows for their individualization from the aorta. The cellular extrusion and release from the aorta take on average 7.5h±1.3h (counted for 10 cells in 10 individuals). Remnant ECs finally complete the remodelling of the aorta by fully recovering its initial diameter and cylindrical geometry (Fig. 1e-f and 5).

## Discussion

In this article, we study and characterise the EHT process at the cellular and tissular level. We quantify the temporal sequences of cells rearrangements, shape modifications, and their coordination with cellular migration events, whilst the DA tissue takes a peculiar tridimensional shape in the trunk region. Thanks to 4D imaging and drug treatment, we have organized the EHT process in two different phases of DA dilation and contraction, and analysed the cellular events occurring.

To decipher the mechanisms that control the DA shape at various stages of EHT, we discuss here why the mechanical properties of the DA endothelium play an important role in the global control of the spatiotemporal organisation of the DA.

In the last century, theoretical methods of classical elasticity have been successful to describe instabilities in metallic pipes and similar mechanical systems that, once subjected to a critical load, change their shape and buckle. More recently, the generalization and adaptation of linear elasticity theory for the needs of biophysical systems has allowed the mechanical description of living matter like lipid membranes, cells or tissues ^24–26^. Moreover, high susceptibility of biological systems, residing near the critical point to the variation of external parameters, is supposed to be often used in nature to control and regulate various processes ^27^

We argue here that mechanical instabilities are involved in the DA shape transition during EHT. More precisely, global variations of aorta shape in the trunk region assisting EHT are the result of instabilities driven by mechanical stresses of various natures acting on the DA. These stresses emerge spontaneously from the DA inhomogeneous growth and its interaction with surrounding tissues rather than from genetically preprogrammed features of embryogenesis. As a matter of fact, the specific diameter modulation in the lower part of the DA appears even when the morphogenetic program of EHT is strongly altered under inhibition of the *runx1* transcription factor ^14^. Moreover, cytoskeleton-blocking drugs only affect the final step of extrusion of HPSC, but not at all the previous phases involving the cellular tissue remodeling and the global shape change of the DA.

The DA is formed by a single monolayer of cells contrary to an adult aorta multilayer complex organization ^14,28^. Therefore, in first approximation, DA can be considered as a thin cylindrical elastic shell. DA is much softer than metallic pipes, and its shape is finally defined by the opposition of different active factors. These factors include: the inner hemodynamic forces, the outer compressions exerted by the surrounding tissues due to embryonic growth and development ^17^ and the in-plane stresses generated in the aorta tissue by ECs shaping and migration as observed in our experiments. The typical timescales for such developmental events are of several hours, thus much longer than the typical time of aortic elastic response to the heart-beat (fractions of seconds), for instance ^29,30^.

Classical mechanics predicts that destabilization of a tubular membrane can be associated with three main modes corresponding to various membrane deformations. These modes are: *Euler buckling* (a tube’s longitudinal axis bends while cross-section remains circular, Fig. 5b), *transverse buckling* (axis of a tube remains straight, whereas its cross-section takes an oval shape Fig. 5c) and *“corrugation instability”* (a tube preserves its rotational symmetry along the main axis, but its radius is modulated along the main axis, Fig. 5d).

Euler buckling of the cylindrical tube can be invoked by the compressive stress along the tube main axis. If the surrounding tissues were absent, the DA stability with respect to this mode would have been determined almost exclusively by the bending rigidity of the tube wall. However, as it is proven by the experiment, long-wave buckling instability is irrelevant for the aorta, because notochord and other tissues surrounding the DA prevent its longitudinal axis from bending (Fig. 1 and 5).

Transverse buckling of the cylindrical shell occurs when transverse isotropic (i.e. possessing the rotational symmetry of the tube) stress in its walls reaches a negative critical value. Stability with respect to transverse buckling is independent on the tension applied along the tube axis and is determined by material constants characterizing the system. This type of mechanical instability also occurs in ordinary rings^31^. Slightly elongated shape of aorta in the dorsal-ventral direction is preserved throughout the whole process and in our opinion this shape is due to an anisotropic compression applied by the muscles located on the sides of the aorta rather than to the transverse buckling instability with spontaneous symmetry breaking.

On the contrary, deformation appearing in the DA prior to HSPCs extrusion has a well-pronounced space periodicity and amplitude (Fig. 1c-e,1h, and Fig. 5) which makes it similar to the corrugation deformation of an axially compressed thin-walled rigid pipe ^32^. Importantly, space period and amplitude of the DA diameter modulation lay beyond the cell diameter and thickness. As it was mentioned before, the blood pressure creates positive stress in DA walls. Its longitudinal component can be calculated as σ=R ΔP/2h, where ΔP is a pressure difference between the interior and the exterior of DA, R and h are DA’s radius and thickness respectively. This positive stress cannot induce corrugation since it makes DA even more stable. However, negative compressive stress (inducing the corrugation in the system) can originate from the difference between the growth rates of the DA and the tissues around it. As it is shown by our data at about 30 h.p.f., this negative stress eventually overpowers the positive one associated with blood pressure and leads to the corrugation of the aorta. This deformation occurs mainly in the ventral part of the DA where the migrating cells converge, and the compressive stress is maximal. Later on, as the zebrafish embryo develops, exit of the cells undergoing EHT decreases the effective stress due to the decrease of the equilibrium surface area of the DA. Consequently, at about 65 h.p.f. the stress drops below the critical value and the tube regains its initial not-deformed shape (Fig. 1f, Fig. 5).

As we show experimentally, chemical perturbation or inhibition of acto-myosin contractility machinery seriously affects only the final event of EHT, i.e. the extrusion of the EHT cell from the aorta endothelium. Very interestingly, we stress that the initial phases of EHT resulting in DA shape distortion still occurs in presence of chemical perturbations. Moreover, the deformation of HSPCs that are preparing to leave the aorta also occurs. What is different is that, in the drug-treated specimens where the actin rings surrounding the cells are absent, many cells do not bend outside (as usual) but do bend inside of the aorta. Based on this observation, we hypothesize that the EHT process is also associated with additional shape instability of individual cells forming the aorta endothelium. We believe that this second instability is also provoked by the stress exerted on the DA tissue, whereas the polymerization of the actin ring and the actin cytoskeleton activity insures the right direction of future HSPC bending and further facilitates its extrusion in the sub-aortic space toward the cardinal vein.

In conclusion, by using 4D fluorescence microscopy, we have characterized qualitatively and quantitatively different dynamical phases of EHT leading to the generation of circulating hematopoietic stem cells from the DA. From the analysis of our observations on wild-type, genetically-modified and chemically-treated zebrafish, we confirmed the important role of the acto-myosin system in EHT single cell final extrusion ^19,33^. Other processes involving the externalisation of a single cell from a cell layer, such as apoptotic extrusion from the epithelial layer of the zebrafish embryo epidermis, confirm that actin/myosin contraction is essential in this process ^34^.

In particular, we profiled a general mechanism based on mechanical instabilities that prepare and support the whole EHT prior to a specific genetic control of the process. Importantly, our interpretation suggests a generic and self-organized mechanism that drives unique collective events of tissue reorganization such as EHT in the development and growth of complex organisms.

Further on, it will be interesting to develop a more precise mathematical model for the description of aorta dynamics and study EHT transition in other model systems. Since the aorta of the zebrafish embryo consists in a limited number of cells, which decreases even more during the EHT process, it will be important to combine an analytical continuum model with a coarse-grained approach allowing for the description of individual cells, similarly to ones used in ^35–37, 38^, but generalized for a curved 2-dimensional surface in 3-dimensional space.

We believe that further studies of EHT will shed light on complex HSPC genesis, a fundamental example of developmental process with important applications in tissue engineering and regenerative therapies, but also on mechanical processes resulting in the development of pathologies. Finally, does mechanics prepare the tissue before genetic reprogramming? It is a debate that is developed in this study in an illustrated way with the example of HSPCs ontogenesis.

## Methods

### Zebrafish husbandry

*Tg*(*kdrl:Has.HRAS-mCherry*) (here cited as *kdrl:caax-mCherry*)^39^, *Tg*(*kdrl:utrophin-CH-GFP*), *Tg*(*Cdh5:Gal4//UAS:lifeact:GFP*)^22^, *Tg*(*kdrl:nls-GFP*)^40^, *Tg*(*flk-1:mV-zGem*)^20^ and *Tg*(*cd41:eGFP*)^23^ were maintained, crossed, raised and staged as described previously ^41, 42^. All animal experiments described in the present study were conducted at the University of Montpellier according to European Union guidelines for handling of laboratory animals (http://ec.europa.eu/environment/chemicals/lab_animals/home_en.htm) and were approved by the Direction Sanitaire et Vétérinaire de l'Hérault and Comité d'Ethique pour lΈxpérimentation Animale under reference CEEA-LR-13007.

### Drug treatments

From 24 h.p.f. onwards, embryos were grown in PTU-containing medium to block pigmentation. Embryos were dechorionated and treated with different drugs after the start of circulation at 26 h.p.f. Drugs were dissolved in DMSO as stock solution and diluted in E3 medium up to 1% DMSO final concentration. Control embryos were subjected to the same concentration of DMSO as treated embryos. Embryos were treated with 2μM Latrunculin B, 0.8μM Blebbistatin, 50μM Rac inhibitor NSC23766 for or 50μM Rho-kinase (ROCK) inhibitor Y-27632. Embryos were kept in drug solution until image acquisition and appropriate concentrations of drugs were added to the mounting medium for time-lapse acquisitions.

### Quantification of HSPC colonization using *Tg*(*cd41:eGFP*)

To quantify cd41+ cells in the CHT, embryos were imaged under a fluorescent binocular scope Zeiss V12 at 100X at 52 h.p.f. and 72 h.p.f. Cd41+ cells were then counted using ImageJ.

### Microscopy

Fluorescence images of transgenic embryos were acquired using Zeiss LSM510 at 20X. Time-lapse imaging was performed using Zeiss LSM510 at 20X or 40X magnification essentially as described^43^. Embryos were anesthetised with tricaine (0.016%) and mounted on a glass petri dish with 0.7 % low melting agarose and covered with standard E3 medium supplemented with tricaine and 1-phenyl-2-thiourea (PTU) (0.003%) to prevent pigment formation. Temperature was maintained at 28°C by placing the dish in a temperature-control chamber during time-lapse acquisitions. Images were analysed using ImageJ and Imaris (Bitplane).

### Aorta diameter measurement

To measure aorta diameter *Tg*(*kdrl:caax-mCherry*) embryos were imaged using Zeiss LSM510 as described above. Aorta diameter was measured manually using ImageJ at 10 different points along the trunk which were then averaged. To show aorta diameter evolution through time, the aorta was measured every 2.5 hours in 5 embryos.

### Cell surface area measurements

Cell surface area was measured using the Surface Contour tool of the Imaris software (Bitplane). Briefly, the contour of a given cell was outlined on the different stacks where the cell was visible using the Click drawing mode. The software then calculated automatically the total cell surface area (μm²). A total of 6 ECs starting to emerge from the aorta floor and 5 ECs on the side of the aorta were analysed in 5 different embryos.

### Statistical analysis

Normal distributions were analysed using Shapiro–Wilk test. Non-Gaussian data were analysed using Wilcoxon or Kruskal–Wallis test, Gaussian with Student’s *t*-test or analysis of variance followed by Holm’s multiple comparison. *P*<0.05 was considered as statistically significant. Statistical analyses were performed using R software.

## Supporting information

Suppl fig et table-Poullet et al

Suppl MovieS1-Poullet et al

Suppl MovieS2-Poullet et al

Suppl MovieS3-Poullet et al

## Acknowledgements

We thank Etienne Lelièvre for his critical reading of the manuscript, A. Sahuquet, C. Chevalier, V. Diakou for their assistance and the MRI facility, N. Abdellaoui for management of zebrafish facility. D. Stainier lab for *Tg*(*Cdh5:Gal4//UAS:lifeact:GFP*), S. Shulte-Merker lab for *Tg*(*kdrl:utrophin-CH-GFP*) and *Tg*(*kdrl:nls-GFP*) and National Bioresource Project Zebrafish for *Tg*(*flk-1:mV-zGem*). This work was supported by the ARC, FRM, ATIP-Avenir fellowships and a fellowship from the Région Languedoc-Roussillon, Chercheur d’Avenir. NP was supported by a fellowship from the ATIP-Avenir, SR and IG are grateful to the RSF grant N 19-12-00032, AP, IG and SR acknowledge NUMEV (AAP-2016-2-025) for financial support. I.G.’s thesis was funded by Campus France (Vernadsky Fellowship) and the France-Russia Cooperation Program, and JT by a fellowship from the MESR and the FRM.

