## Supplementary material for "Mechanical instabilities of aorta drive blood stem cell production: a live study": Suppl fig et table-Poullet et al

**Supplementary Figures legends.**

**Supplementary Figure 1.**

**a-d**. Single z-stacks of time-lapse imaging of *Tg(kdrl:utrophin-CH-GFP)/Tg(kdrl:caax-mCherry)* embryo during EHT after cytoskeleton drug treatments. After Latrunculin B (**a**), Rac inhibitor (**b**), Blebbistatin (**c**) or ROCK inhibitor (**d**) treatment, the actin ring is no longer visible and some EHT events occur towards the aorta lumen. Scale bar: 10 µm.

### Supplementary Figure 1

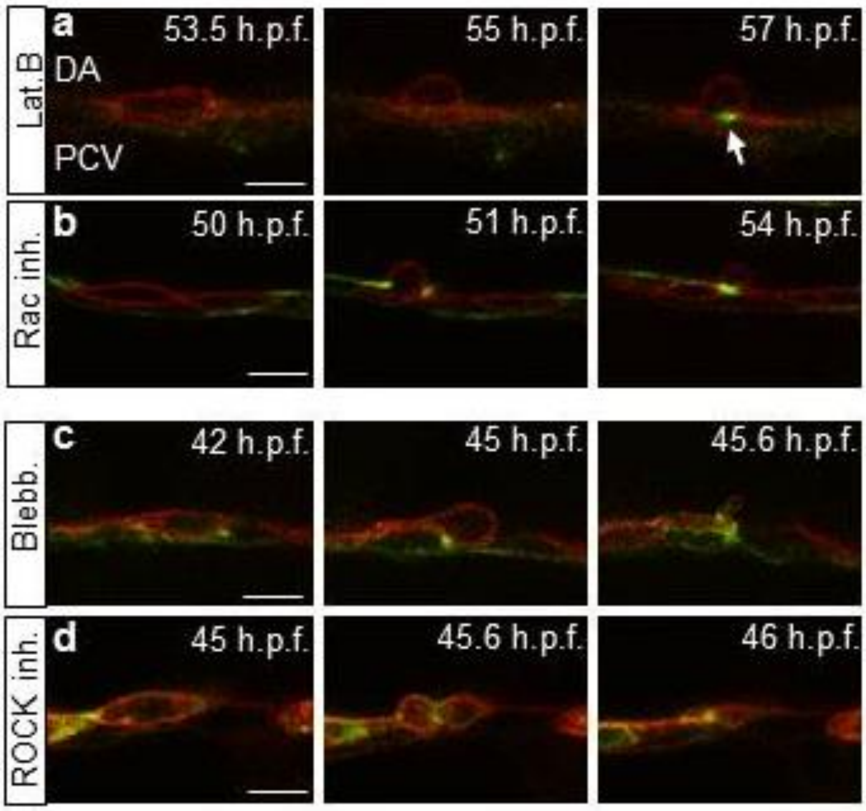

**Supplementary Table 1**. Number of HSPCs (cd41+ cells) colonizing the Caudal Hematopoietic Tissue (CHT) between 52 and 72 h.p.f. in control and cytoskeleton drug-treated embryos.

| Compound | Control | Treated | p-value |
|---|---|---|---|
| LatrunculinB | 47.8±3.3 | 25.2±2.0 | 3.7E-5 |
| Blebbistatin | 84.6±4.3 | 69.4±3.8 | 0.01 |
| Rac inhibitor | 89.6±1.2 | 51.0±10.2 | 0.03 |

**Supplementary Movie S1**. Time lapse confocal fluorescence imaging of an embryo from 30 to 65 h.p.f.: diameter temporal evolution.

This movie goes together with Figure 1. It shows a time lapse confocal fluorescence imaging of a *Tg(kdrl:caax-mCherry)* embryo from 30 to 65 h.p.f. This movie shows a maximum projection from 40 z-stacks spaced by 1µm. DA diameter first increases between 30 and 45 h.p.f., and then decreases between 45 and 65 h.p.f. Decrease of DA diameter corresponds to the peak of EHT events.

**Supplementary Movie S2**. Time lapse confocal fluorescence imaging of an embryo from 42 to 60 h.p.f.: dynamics of endothelial cell junctions during EHT.

This movie shows a time lapse confocal fluorescence imaging of a *Tg(kdrl:utrophin-CH-GFP)* embryo from 42 to 60 h.p.f. This movie shows a maximum projection from 40 z-stacks spaced by 1µm. It allows the visualisation of the dynamics of EC junctions during EHT.

**Supplementary Movie S3**. Time lapse confocal fluorescence imaging of an embryo from 45 to 60 h.p.f.: formation and closure of the actin ring surrounding EHT cells.

This movie goes together with **Figure 4a**, **e-g**. It shows a time lapse confocal fluorescence imaging of a *Tg(kdrl:utrophin-CH-GFP)/Tg(kdrl:caax-mCherry)* embryo from 45 to 60 h.p.f. This movie shows a 3D view obtained with Imaris software, allowing visualisation of the AGM from a horizontal reconstructed view. Formation and closure of the actin ring surrounding EHT cells is observable in cells 1, 2, 3.
